# The *Toxoplasma* Oxygen Sensing Protein, TgPhyA, Is Required for Resistance to Interferon-gamma Mediated Nutritional Immunity

**DOI:** 10.1101/2023.05.16.540931

**Authors:** Charlotte Cordonnier, Msano Mandalasi, Jason Gigley, Elizabeth A. Wohlfert, Christopher M. West, Ira J Blader

## Abstract

As *Toxoplasma gondii* disseminates through its host, the parasite must sense and adapt to its environment and scavenge nutrients. Oxygen (O_2_) is one such environmental factor and cytoplasmic prolyl 4-hydroxylases (PHDs) are evolutionarily conserved O_2_ cellular sensing proteins that regulate responses to changes in O_2_ availability. *Toxoplasma* expresses two PHDs. One of them, TgPHYa hydroxylates SKP1, a subunit of the SCF-E3 ubiquitin ligase complex. *In vitro,* TgPHYa is important for growth at low O_2_ levels. However, studies have yet to examine the role that TgPHYa or any other pathogen encoded PHD plays in virulence and disease. Using a type II ME49 *Toxoplasma* TgPHYa knockout, we report that TgPHYa is important for Toxoplasma virulence and brain cyst formation in mice. We further find that while TgPHYa mutant parasites can establish an infection in the gut they are unable to efficiently disseminate to peripheral tissues because the mutant parasites are unable to survive within recruited immune cells. Since this phenotype abrogated in IFNγ knockout mice, we studied how TgPHYa mediates survival in IFNγ-treated cells. We find that TgPHYa is not required for release of parasite-encoded effectors into host cells that neutralize anti-parasitic processes induced by IFNγ. In contrast, we find that TgPHYa is required for the parasite to scavenge tryptophan, which is an amino acid whose levels are decreased after IFNγ upregulates the tryptophan-catabolizing enzyme, indoleamine dioxygenase (IDO). We further find that relative to wild-type mice that IDO knockout mice display increased morbidity when infected with TgPHYa knockout parasites. Together, these data identify the first parasite mechanism for evading IFNγ-induced nutritional immunity and highlight a novel role that oxygen sensing proteins play in pathogen growth and virulence.

## Introduction

*Toxoplasma gondii* is an intracellular apicomplexan parasite that chronically infects up to one-third of the world’s population [1]. This opportunistic pathogen causes toxoplasmosis, a disease that can lead to life-threatening disease in immunocompromised individuals and developing fetuses. Infection in humans follows consumption of meat harboring bradyzoite-containing tissue cysts or water and produce contaminated with sporozoite-containing oocysts [2]. Gastric enzymes rupture cysts or oocysts and the released parasites go on to infect the small intestine. At this juncture, the parasites convert into replicative tachyzoites and trigger an immune response that results in recruitment of innate immune cells such as inflammatory monocytes, innate lymphoid cells, and neutrophils. These cells are in turn infected and used to disseminate to peripheral tissues where tachyzoites differentiate into bradyzoites and form tissue cysts that can persist for the host’s lifetime [3].

As *Toxoplasma* traffics through a host it encounters diverse environments that vary in essential nutrient availability. One such nutrient is O_2_, which all cells must sense and respond to changes in its levels to optimize growth and viability [4]. Cytoplasmic prolyl 4-hydroxylases (PHDs) are dioxygenases that use α-ketoglutarate and O_2_ as substrates and function as the key cellular O_2_ sensors [5]. PHDs hydroxylate proline residues in substrates and in metazoans the best-recognized PHD substrate is the α subunit of the hypoxia-inducible transcription factor (HIFα). Prolyl-hydroxylated HIFα is ubiquitinated by the Von Hippel-Lindau ubiquitin ligase complex, which targets it for proteasomal degradation [6]. Under hypoxic conditions, HIFα is stabilized and able to regulate the expression of genes important for growth and survival at low O_2_. While HIFα is not conserved in protist genomes, PHDs are, indicating that protists likely respond differently to changes in O_2_ availability [7, 8]. The best characterized protozoan PHD is DdPhyA from *Dictyostelium discoideum* that modifies a proline in DdSkp1, which is an adaptor protein in the SCF (Skp1/Cullin/F-box protein) E3 polyubiquitin ligase complex [9]. The resulting hydroxyproline is subsequently modified by a series of glycosyltransferases that alters the binding affinity of SKP1 towards different F-Box proteins and thus differential protein ubiquitination by the SCF complex [10–12]. Two PHDs have been identified in *Toxoplasma* and one of these, TgPHYa also modifies SKP1 [13, 14]. While TgPHYa is not essential, TgPHYa knockouts in type 1 strain parasites display reduced *in vitro* growth at low O_2_ [15]. But TgPHYa’s function *in vivo* remains to be determined.

Interferon gamma (IFNγ) is a cytokine critically required for host resistance to *Toxoplasma* [16, 17]. It acts by triggering the expression of IFNγ-responsive genes whose products are involved in oxygen radical generation and degradation of the parasitophorous vacuole membrane via the activity of immunity-related guanosine triphosphatases (IRGs) and guanylate binding proteins (GBPs) [18]. In addition, IFNγ induces the expression of Indoleamine-pyrrole 2,3-dioxygenase (IDO) that catabolizes tryptophan, which *Toxoplasma* must scavenge from its host [19]. The importance of IDO in nutritional immunity to *Toxoplasma* has remained enigmatic as it is important for resistance in some human cells but appears to be dispensable in mice [19–21]. The most likely reason for this is that unlike humans, mice express two IDO isoforms and thus it remains unknown whether tryptophan scavenging/utilization remains a viable anti-parasitic drug target [21].

Here, we report that TgPHYa deletion in a cystogenic strain leads to decreased virulence in both per oral and intraperitoneal infection models. Furthermore, TgPHYa is required for resistance to IFNγ-mediated killing and does so by evading IFNγ-inhibition of tryptophan utilization. Significantly, TgPHYa knockout parasites display increased virulence in IDO1 knockout mice. Taken together, these data reveal that a parasite-encoded cytoplasmically localized prolyl hydroxylase mediates *Toxoplasma* evasion of a protective host nutritional immune response pathway.

## Results

### Generation of TgPHYa Knockout in the Type II ME49 Strain

To assess a role for TgPHYa during *in vivo* infection, CRISPR was used to generate a TgPHYa knockout mutant in a type II *Toxoplasma* strain that also expresses red fluorescent protein (RFP) (TgPHYaKO_II_). PCR analysis indicated that TgPHYa was properly targeted by the CRISPR construct and that the TgPHYa gene was disrupted (**Fig. 1A&B**). Because TgPHYa antisera is lacking, we assessed whether the gene disruption affected TgPHYa protein by analyzing TgSKP1 mobility by SDS-PAGE. We observed that the apparent molecular weight of TgSKP1 was reduced in TgPHYaKO_II_ parasites indicating decreased TgSKP1 prolyl hydroxylation/glycosylation (**Fig. 1C**) [15]. Similar to the type I strain TgPHYa knockout [15], *in vitro* TgPHYaKO_II_ growth was slightly decreased at 21% O_2_ **(see Fig. 8A&B).**

**Figure 1.**
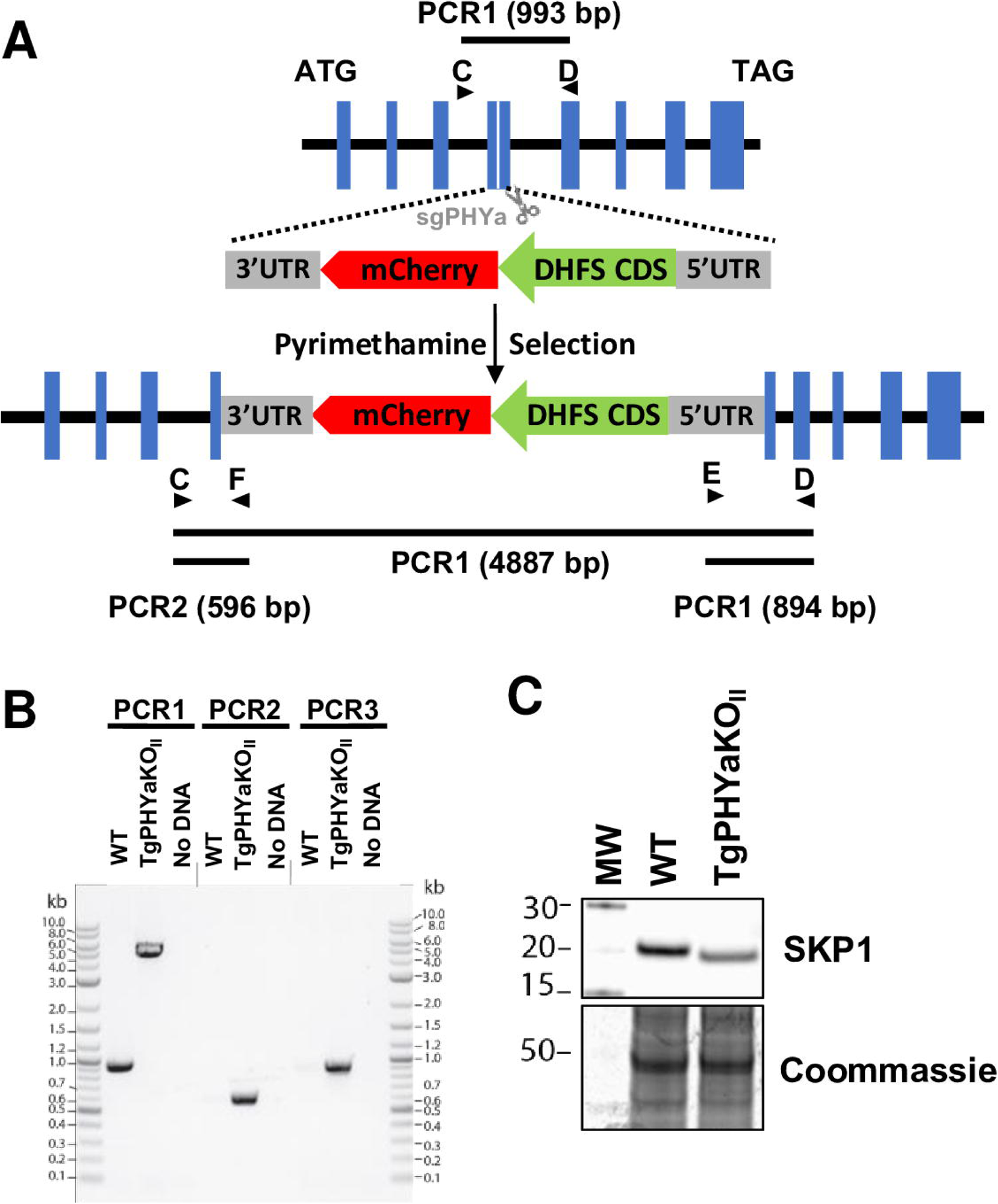
TgPHYa Knockout in Type II Strain Parasites. **(A)** Scheme for disrupting the TgPHYa locus by CRISPR. **(B).** Genomic DNA from wild-type and TgPHYaKO_II_ parasites were analyzed by PCR using the primers depicted in **(A)**. **(C)** Western blot analysis to detect SKP1 in lysates from the indicated strains. Note the increased mobility of the SKP1 in the TgPHYaKO_II_ lysate indicating is loss of prolyl hydroxylation/glycosylation.

### TgPHYaKO_II_ Displays Reduced Virulence and Decreased Numbers of Brain Cysts

C57BL/6J mice were infected by oral gavage with 50 or 100 wild-type (WT) or TgPHYaKO_II_ tissue cysts and survival and body weight were monitored for 30 days. All mice infected with 50 TgPHYaKO_II_ cysts survived up to 30 days post infection, while 40% of mice infected with the parental WT strain died (**Fig. 2A**). Moreover, all mice infected with 100 WT cysts succumbed during the acute stage of the infection (within 8 days post infection) while <40% of the mice succumbed when infected with 100 TgPHYaKO_II_ cysts (**Fig. 2A**). Similar to decreased mortality, mice infected with the TgPHYa knockout lost less weight than those infected with wild-type parasites and their weight loss rebounded whereas weight loss was sustained in the surviving wild-type infected mice (**Fig. 2B**).

**Figure 2.**
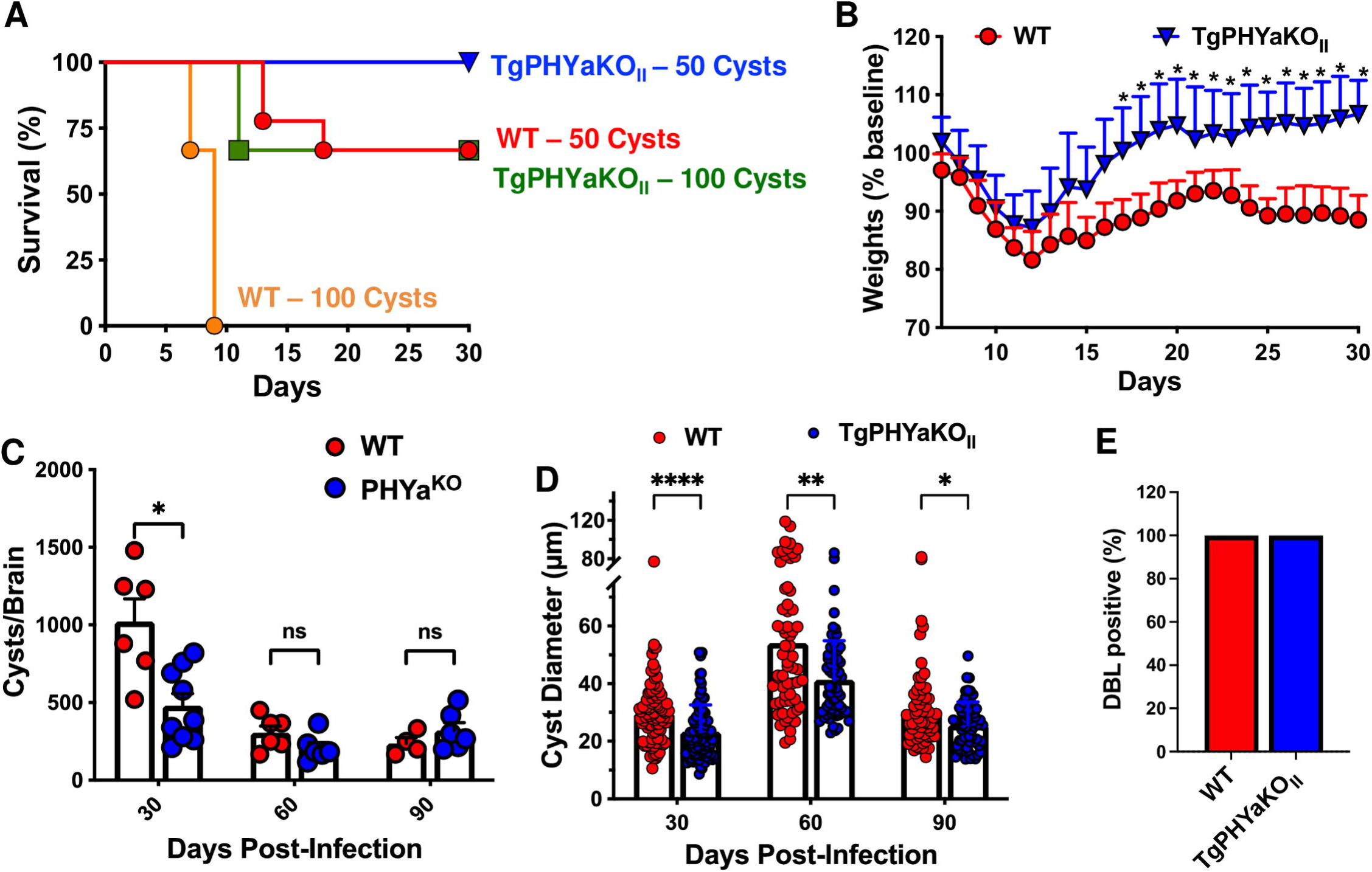
C57BL/6 mice are less susceptible to TgPHYaKO_II_ strain infection correlating with fewer number of brain cysts. **(A)** Survival of C57BL/6J mice infected orally with 50 or 100 cysts of the indicated strains. Cumulative data from 3 independent trials and 9 mice for each strain and dose. Curves were compared by log-rank survival analysis of Kaplan-Meier curves, 50 cysts p=0.0649, 100 cysts \*\*\**P<*0.0001. **(B)** Weight loss was monitored in mice infected with 50 cysts. Shown are cumulative data from 3 independent trials (mean ± SD \**P<*0.05, multiple Student’s *t* test with Holm-Sidake correction). Brain cyst burden **(C)** and diameter **(D)** were calculated from mice 30, 60 or 90 days after infection. Shown are cumulative data from 3 independent trials (mean ± SD \**P<*0.05, multiple Student’s *t* test with Holm-Sidake correction). **(E)** *In vitro* cyst development was assessed by Dolichos lectin staining of parasites grown in HFFs for 5 days in in pH 8.2 medium. Shown are percentage of lectin^+^ vacuoles from 3 independent assays (50 randomly selected vacuoles were counted for each sample).

Next, brain cyst burdens were enumerated in mice 30-, 60-, or 90-days post infection (dpi) with 50 cysts. We found that at 30 dpi there was a statistically significant decrease in numbers of cysts in brains of TgPHYaKO_II_ infected mice (**Fig. 2C**). But, at later time points differences were not noted. Brain cyst sizes were also analyzed and at each time point the TgPHYaKO_II_ cysts were significantly smaller than the parental strain cysts (**Fig. 2D**). To test whether TgPHYaKO_II_ brain cyst phenotypes were a result of the prolyl hydroxylase functioning in cyst formation, cyst formation was examined in parasites exposed to alkaline medium (pH 8.2) at ambient CO_2_ [22]. After 5 days of induction, cysts were examined using DBA-FITC and no differences in the ability for TgPHYaKO_II_ to form *in vitro* cysts were noted (**Fig. 2E**). Together, these data indicate that TgPHYa is important for virulence and brain cyst development.

### TgPHYa is Important for Parasite Dissemination

One possible explanation for decreased numbers of TgPHYaKO_II_ brain cysts at 30 dpi is delayed dissemination from the gut to the brain and other tissues. To examine this, we first assessed a role for TgPHYa in establishing an infection in the gut by gavage infecting mice with 50 WT or TgPHYaKO_II_ cysts and 7 days later enumerating parasite burdens in different segments of the small intestine. No significant differences in parasite numbers were observed throughout the intestine (**Fig. 3A**). We also assessed intestinal inflammation by histological analysis at 7 dpi (**Fig. 3B**) and 12 dpi (**Fig. 3C**) and no significant differences were noted at either time point.

**Figure 3.**
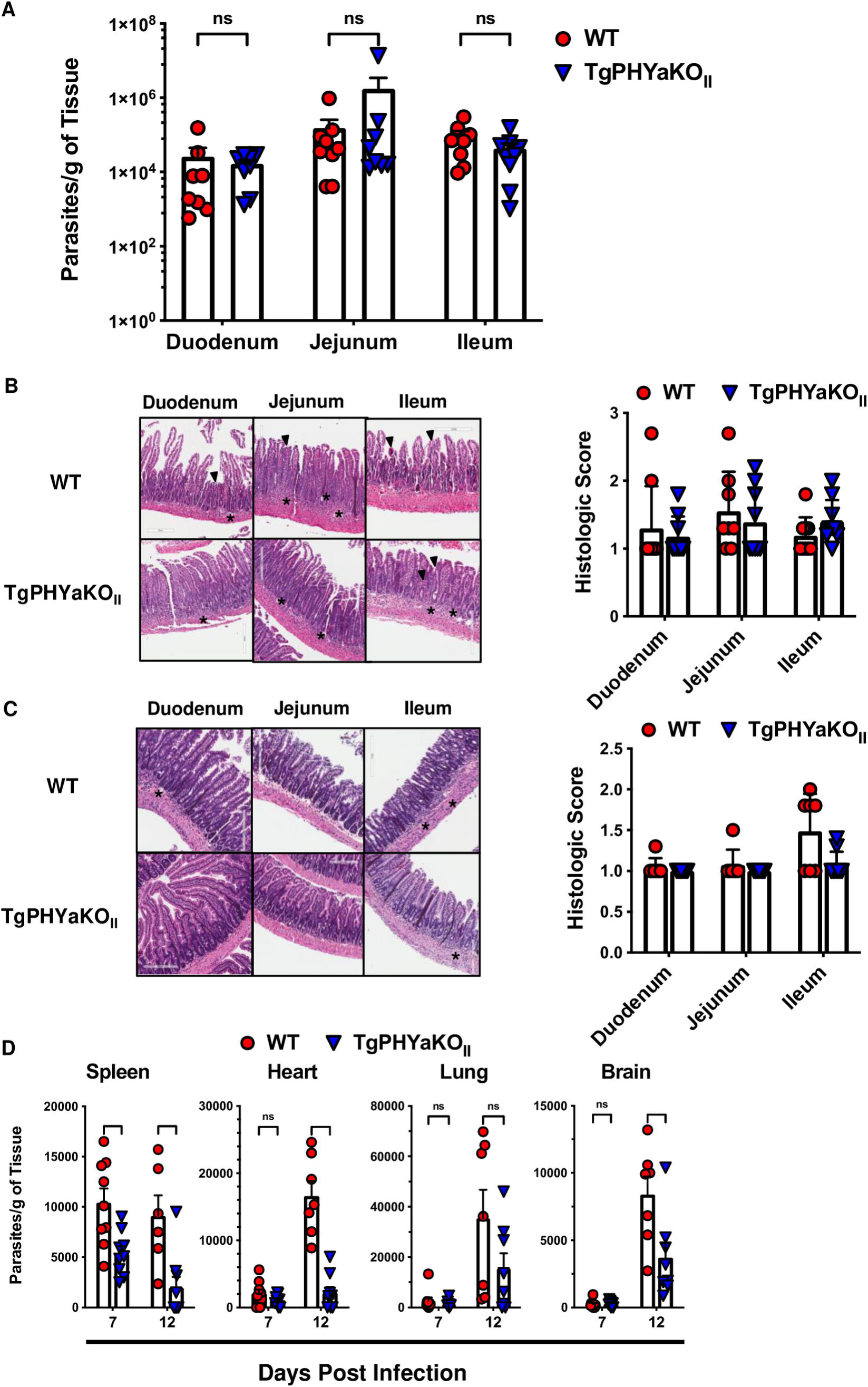
TgPHYaKO_II_ is not required for colonization of intestinal epithelium but is important for parasite dissemination to peripheral tissues. **(A)** ME49 WT or TgPHYaKO_II_ burdens were analyzed by quantitative RT-PCR in the indicated intestinal sections 7 days following gavage infection with 50 cysts. Shown are means ± SD of 3 independent experiments with 3 mice per experiment (*ns= not significant*, using one-way ANOVA). **(B,C)** H&E-stained small intestinal sections from mice gavage infected with 50 WT or TgPHYaKO_II_ cysts on day 7 **(B)** or day 12 **(C)** post infection. Asterisks highlight inflammatory infiltrates in the villi and lamina propria, and arrowheads highlight destruction of villi. Shown are representative images and plots represent mean ± SD of histological scores of 50 randomly selected section per mouse. **(D)** Parasite burdens were determined by qPCR of genomic DNA isolated from spleens, lungs, hearts and brains of mice gavage infected with 50 of ME49 WT or TgPHYaKO_II_ cysts. Data shown are mean ± SD from a total of 7 to 9 mice from 3 independent experiments. (*ns= not significant*, \*\**P<*0.01, \*\*\**P<*0.001, using one-way ANOVA).

Next, parasite burdens in peripheral tissues at 7 and 12 dpi were determined by quantitative real time PCR. Except for the spleen, there were no remarkable differences in parasite burdens in the organs on day 7 of infection (**Fig. 3D**). In contrast, significantly fewer TgPHYaKO_II_ parasites were observed in the spleens, hearts, and brains of the infected mice at 12 dpi. Parasite burdens in lungs were also lower although this difference was not statistically significant. These data therefore indicate that while TgPHYaKO_II_ parasites can establish an infection in the intestine, they are unable to disseminate efficiently to peripheral tissues.

### TgPHYaKO_II_ Burdens are Reduced in Inflammatory Monocytes and Neutrophils

*Toxoplasma* dissemination within and from the gut is dependent on parasite infection of and survival within inflammatory monocytes, neutrophils, and CD103^+^ dendritic cells [23]. To assess whether TgPHYa was required for *Toxoplasma* interactions with those cells, we first examined by flow cytometry the numbers of each cell type in intestines of mice 7 days after they were gavage-infected with 50 WT or TgPHYaKO_II_ tissue cysts. Relative to WT infected mice, we observed lower numbers of neutrophils but not inflammatory monocytes nor dendritic cells in the small intestines of TgPHYaKO_II_ infected mice (**Fig. 4A-B**).

**Figure 4.**
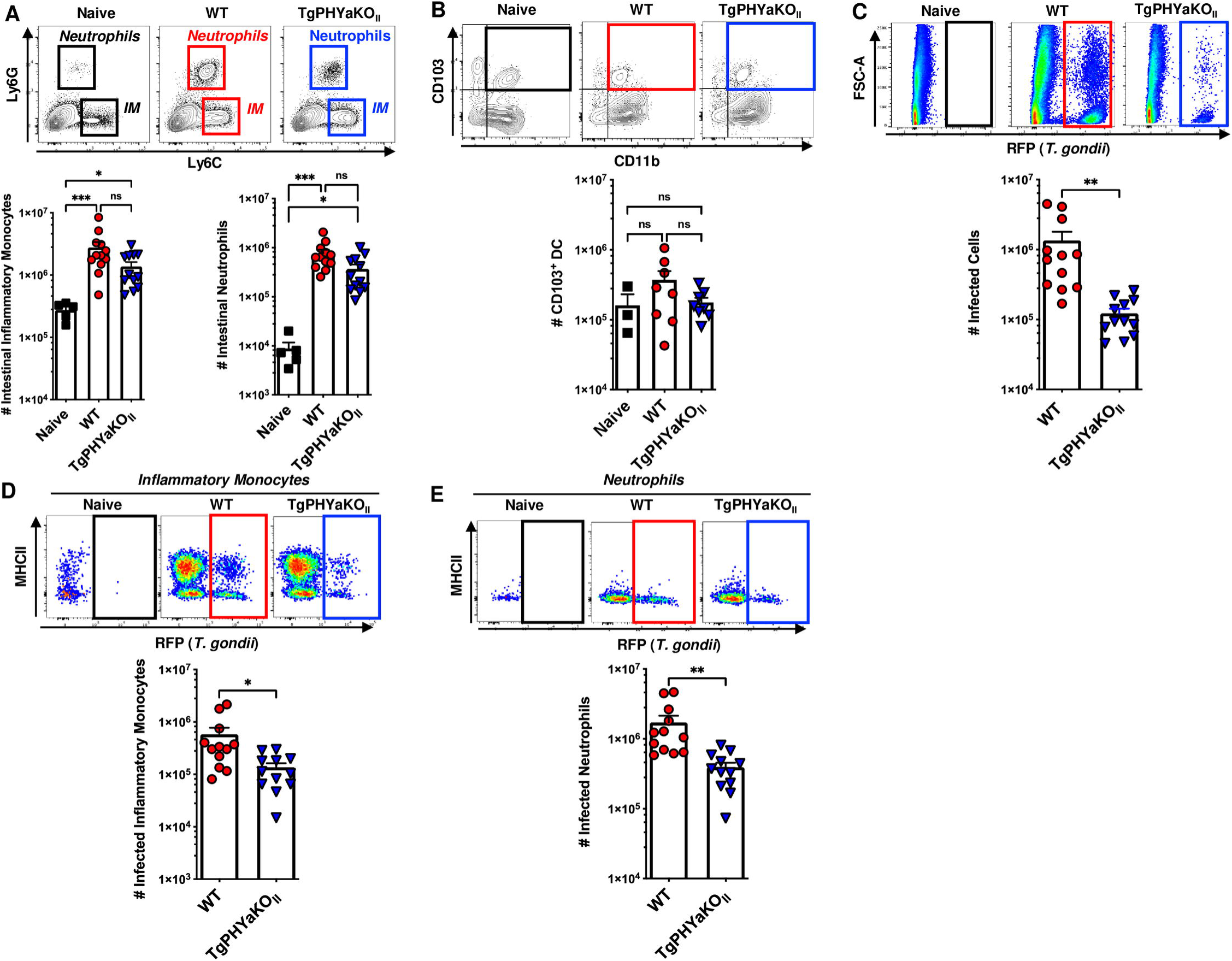
TgPHYa Is Important for Survival in Inflammatory Cells within the Small Intestine. Small intestine lamina propria from uninfected mice and mice gavage-infected with 50 WT or TgPHYaKO_II_ tissue cysts were harvested 7 dpi and analyzed by flow cytometry. Cells were gated on live/CD45^+^/CD11b^+^ cells to identify **(A)** inflammatory monocytes (IM; Ly6C^hi^Ly6G^low^) and neutrophils (Ly6G^hi^Ly6C^int^), and **(B)** CD11b^+^CD103^+^ dendritic cells. (**C**) Parasite burdens were determined by assessing RFP expression of live gated cells. **(D&E)** Numbers of parasite-infected inflammatory monocytes and neutrophils, respectively, were determined by flow cytometry. Shown are representative flow cytometry plots. Graphs represent pooled analyses (mean ± SEM, pooled from five independent experiments); *ns= not significant*, \**P<*0.05, \*\**P<*0.001, \*\*\**P<*0.0001, using unpaired Student *t* test.

The infection status of these cells was assessed by flow cytometric detection of parasite-expressed red fluorescent protein (RFP). First, we compared total numbers of RFP^+^ cells and found significantly decreased numbers of TgPHYaKO_II_-infected cells (**Fig. 4C**). Using the same gating strategies as described above to identify cell populations, no differences in numbers of infected CD11b^+^CD103^+^ dendritic cells (data not shown) were found whereas significantly fewer TgPHYaKO_II_-infected inflammatory monocytes and neutrophils were noted (**Fig. 4D-E**).

To determine if these results were specific to the intestinal environment, we intraperitoneally infected mice with 10^4^ WT or TgPHYaKO_II_ tachyzoites. After 7 days, peritoneal exudates were collected and total numbers and infection status of recruited immune cells were determined. Lower numbers of neutrophils but not inflammatory monocytes were present in peritoneal exudates from TgPHYaKO_II_-infected mice (**Fig. 5A**). As observed in the lamina propria, there were significantly fewer cells infected with TgPHYaKO_II_ parasites (**Fig. 5B**). And, the TgPHYaKO_II_-infected mice had significantly fewer infected inflammatory monocytes and neutrophils (**Fig. 5C-D**). Finally, virulence was assessed by IP infecting mice with high dose of 10^6^ WT or TgPHYaKO_II_ tachyzoites for 7 days. Survival of mice IP infected with TgPHYaKO_II_ parasites was dramatically enhanced as compared to mice infected with the same number of WT parasites (**Fig. 5E)**. Taken together, these data indicate that TgPHYa is important for *Toxoplasma* virulence.

**Figure 5.**
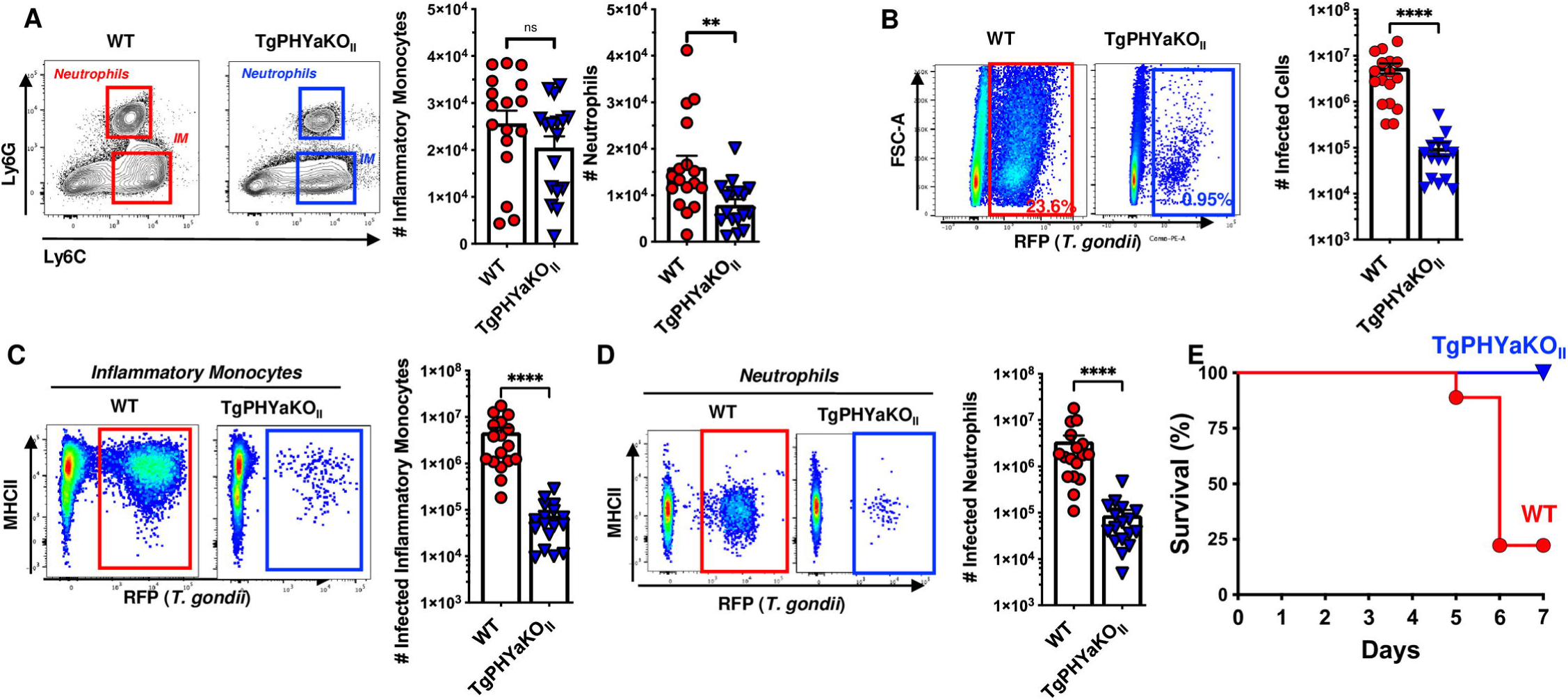
TgPHYaKO_II_ is Important for Survival within the Intraperitoneal Cavity. Mice were intraperitoneally infected with 10^4^ WT or TgPHYaKO_II_ tachyzoites and 7 days later the peritoneal exudate was harvested and processed for flow cytometry. **(A)** Live/CD45^+^/CD11b^+^ gated cells were stained to identify inflammatory monocytes (IM; Ly6C^hi^Ly6G^low^) and neutrophils (Ly6G^hi^Ly6C^int^). (**B**) Parasite burdens were determined by assessing RFP expression of live gated cells. **(C&D)** Numbers of parasite-infected inflammatory monocytes and neutrophils, respectively, were determined by flow cytometry. **(E)** Kaplan-Meier curve showing survival of mice IP infected with 10^6^ WT or TgPHYaKO_II_ tachyzoites. Cumulative data from 3 independent experiments (*n* = 9 total for each strain).

### TgPHYa Is Required For Resistance to IFNγ

Because IFNγ is critically required for host resistance to *Toxoplasma* [24], survival of WT and IFNγ^-/-^ mice gavage-infected with 50 ME49 WT or TgPHYaKO_II_ was monitored for 30 days. Infection with either parasite strain led to rapid death of IFNγ^-/-^ mice (**Fig. 6A)**. Loss of IFNγ also restored parasite growth in peritoneal exudate cells harvested from IP infected mice (**Fig. 6B**). We next compared the peritoneal immune cell infiltrate between WT and IFNγ^-/-^ mice and found significantly higher numbers of inflammatory monocytes in IFNγ^-/-^ mice infected with TgPHYaKO_II_. Moreover, loss of IFNγ led to increased recruitment of neutrophils in mice infected with either WT or TgPHYaKO_II_ parasites (**Fig. 6C**). Finally, we found that IFNγ restricted growth of TgPHYaKO_II_ parasites in both inflammatory monocytes and neutrophils (**Fig. 6D-E**).

**Figure 6.**
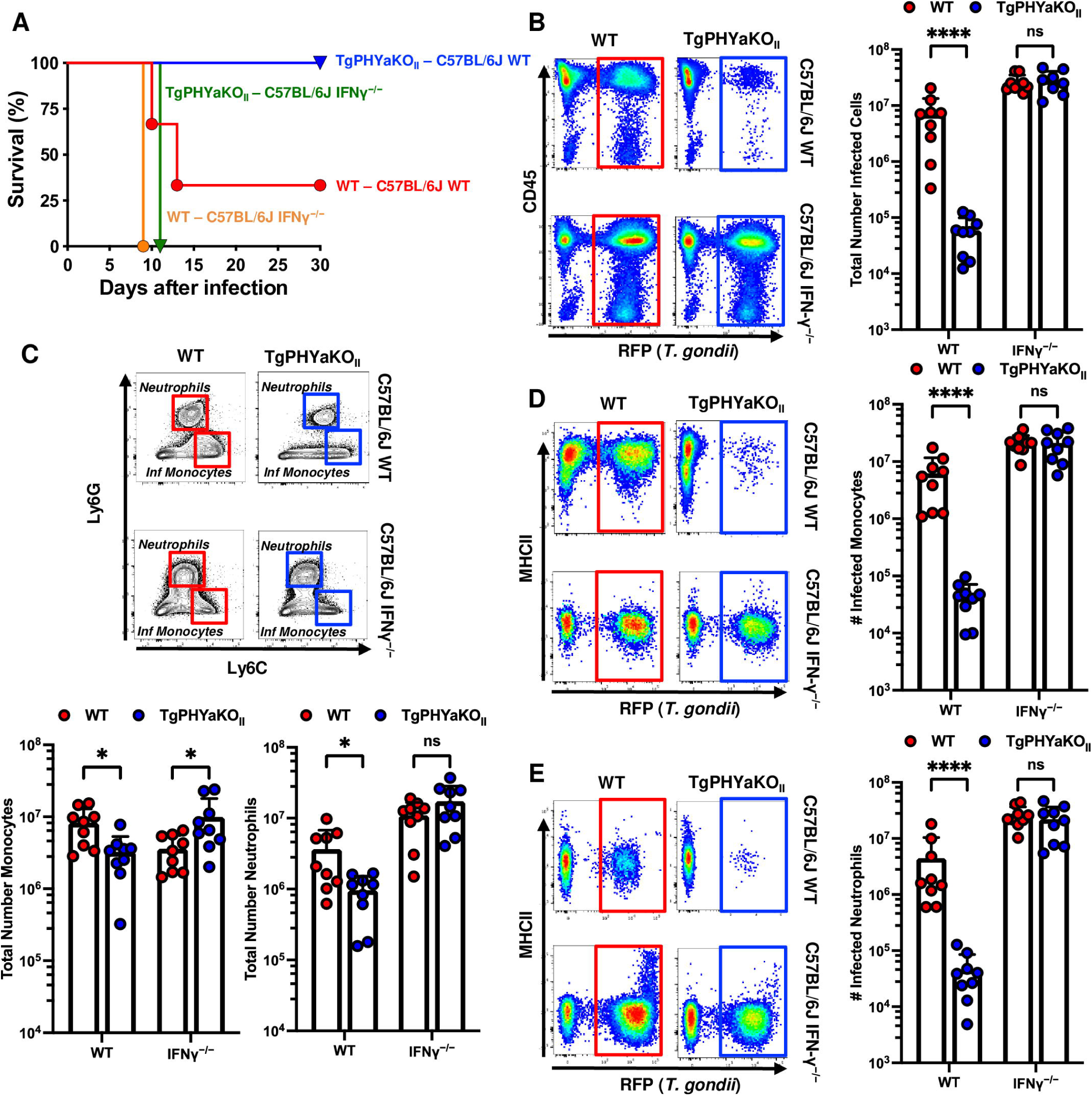
TgPHYa Is Required for Parasite Resistance to IFNγ. **(A)** Kaplan-Meier survival curves of WT and IFNγ^-/-^ mice infected orally with 50 wild-type or TgPHYaKO_II_ cysts. Cumulative data from 3 independent experiments (*n* = 9 total for each strain). **(B-E)** WT and IFNγ^-/-^ mice were intraperitoneally infected with 10^4^ wildtype or TgPHYaKO_II_ tachyzoites. Mice were euthanized after 7 days and peritoneal cavity cells were analyzed by flow cytometry to enumerate numbers of parasite-infected cells **(B)**, inflammatory monocytes **(C)** neutrophils **(D)**, and their infection status **(E**). Shown are representative flow cytometry plots. Graphs represent pooled analyses (mean ± SEM, n = 9 from three independent experiments). *ns= not significant*, \**P<*0.05, \*\**P<*0.001, \*\*\**P<*0.0001, using unpaired Student *t* test.

CCR2^+^ inflammatory monocytes are required for host resistance to *Toxoplasma* and CCR2 deficient mice die during the acute phase of the infection [25, 26]. CCR2^-/-^ mice infected with TgPHYaKO_II_ parasites survived although, relative to wild-type mice infected with TgPHYaKO_II_ parasites, they demonstrated increased weight loss and cyst numbers (**Fig. S1A-C**). Despite an absence of inflammatory monocytes, significantly fewer TgPHYaKO_II_ parasites were found in both wild-type and CCR2 deficient mice (**Fig. S1D**), which was most likely due to elevated (though not statistically greater (p = 0.065) levels of IFNγ between the mouse populations (**Fig. S1E**).

Given the early timing for TgPHYaKO_II_ susceptibility to IFNγ-dependent killing in innate immune cells, we evaluated the importance of Natural Killer (NK) cells as an innate cell source for IFNγ [27]. Thus, NK-cell depleted mice (or those treated with irrelevant IgG) were IP infected. The peritoneal exudate was collected 5 days later and we found that numbers of infected cells were increased in NK-cell depleted mice infected with both WT and TgPHYaKO_II_ strains (**Fig. S2**). Together, these data indicate that a parasite-encoded cytoplasmically localized prolyl hydroxylase, TgPHYa, is required for *Toxoplasma* to escape IFNγ mediated death.

### *Toxoplasma* Growth and Virulence is Dependent on TgPHYa Expression Levels

Two TgPHYaKO_II_ complementation constructs were generated in which the TgPHYa ORF with a 3X-hemagluttin (HA) C-terminal epitope tag was cloned downstream of either the strong *TUB1* promoter (TgPHYaKO_II_:PHYa^TUB^) or its endogenous promoter (1000 bp upstream of its predicted transcription started site; TgPHYaKO_II_:PHYa^PHYa^) (**Fig. S3A&B**). TgPHYaKO_II_:PHYa^TUB^ protein levels were readily detected by Western blotting while TgPHYaKO_II_:PHYa^PHYa^ was not detectable, even by immunoprecipitation (**Fig. 7A**). To assess transgene function, SKP1 mobility was assessed by Western blotting and in both strains SKP1 was found to migrate at a higher molecular weight than it did in TgPHYaKO_II_ lysates indicating that SKP1 was modified as expected (**Fig. 7A**). We also noted that SKP1 protein levels were reduced in the knockout strain suggesting that TgPHYa not only modifies SKP1 but may also affect its abundance.

**Figure 7.**
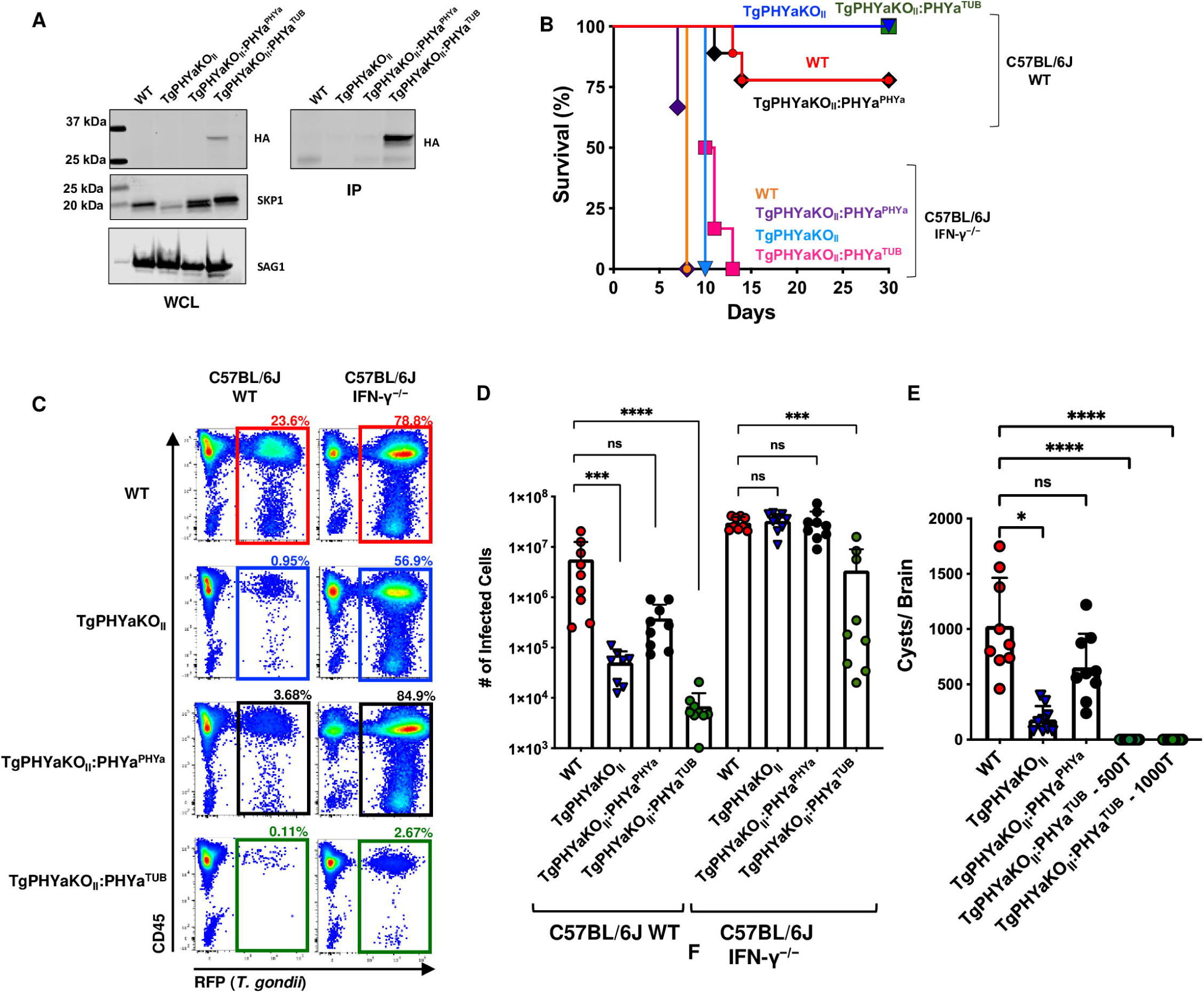
TgPHYa Overexpression of Reduces Virulence. **(A)** Whole cell lysates (WCL) from the indicated strains were either Western blotted to detect SKP1, HA-tagged TgPHYa or SAG1 (as a loading control) or immunoprecipitated (IP) using anti-HA to detect HA-tagged TgPHYa. **(B)** Kaplan-Meier curve showing survival of C57BL/6J WT and IFNγ^-/-^ mice intraperitoneally infected with 10^4^ tachyzoites of the indicated strains. Cumulative data from 3 independent experiments (*n* = 9 total for each strain). **(C&D)** C57BL/6J WT and IFNγ^-/-^ mice were intraperitoneally infected with 10^4^ tachyzoites of the indicated strains. ME49 WT, TgPHYaKO_II_, TgPHYaKO_II_:PHYa^PHYa^ and TgPHYaKO_II_:PHYa^TUB^ parasites. After 7 days infection, mice were euthanized and infection status of peritonealexudate analyzed by flow cytometry. Shown are mean ± SEM, n = 9, pooled from three independent experiments, *ns= not significant*, \**P<*0.05, \*\**P<*0.001, \*\*\**P<*0.001, using unpaired Student *t* test). **(E)** Cysts burdens were determined in brains of mice 30 days after intraperitoneal infection with 500 tachyzoites of the indicated strain or 1000 TgPHYaKO_II_:PHYa^TUB^. Shown are mean ± SEM, n = 9, pooled from three independent experiments, *ns= not significant*, \**P<*0.05, \*\*\*\**P<*0.0001, using unpaired Student *t* test.

To assess complementation of TgPHYaKO_II_ phenotypes, we first examined *in vitro* growth. We observed that the TgPHYaKO_II_ growth defect was restored in TgPHYaKO_II_:PHYa^PHYa^ but not TgPHYaKO_II_:PHYa^TUB^ parasites (**Fig. 8A; compare no treatment samples**). We assessed the importance of TgPHYa expression levels on virulence by first IP infecting mice with 10^4^ parasites of each strain and found that TgPHYaKO_II_:PHYa^PHYa^ was as virulent as WT strain parasites and TgPHYaKO_II_:PHYa^TUB^ was significantly less virulent **(Fig. 7B)** even when mice were IP infected with 10^6^ tachyzoites (**Fig. S4A&B).** Similarly, parasite growth within peritoneal cells was partially restored by TgPHYaKO_II_:PHYa^PHYa^ but not TgPHYaKO_II_:PHYa^TUB^ **(Figs. 7C&D)**. However, both strains were similarly virulent in IFNγ^-/-^ mice **(Fig. 7B)**.

**Figure 8.**
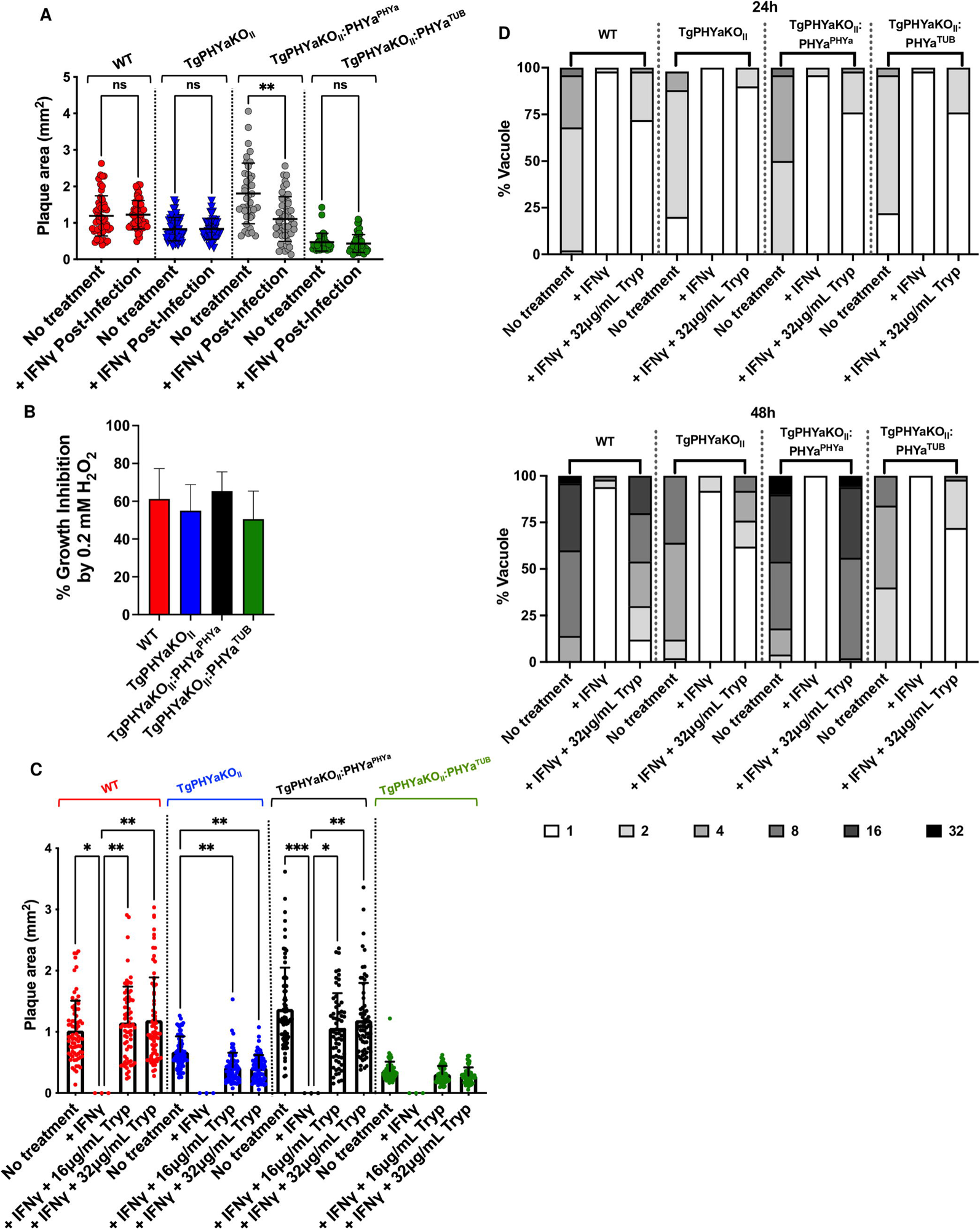
TgPHYa is Important for Tryptophan Utilization in IFNγ-Treated Cells. **(A)** *Toxoplasma*-infected HFFs were mock or IFNγ treated 24 hpi and 12 days later the monolayers were fixed and numbers and sizes of plaques determined (mean ± SEM, pooled from three independent experiments; *ns= not significant*, \*\**P<*0.01 using a multiple comparison 2way ANOVA test). **(B)** Equal number of parasites were added to HFFs and grown in the presence of 0.2 mM H_2_O_2_. After 12 days, monolayers were fixed, numbers of plaques were counted, and results are expressed as the percentage of plaques formed relative to each strain grown without H_2_O_2_. **(C)** HFFs were pretreated with IFNγ (1U/mL) for 24h and then infected with tachyzoites of the indicated strain. The cells were grown for 12 days in the presence of increasing tryptophan concentrations and then the monolayers were fixed and plaque sizes enumerated. **(D)** Bone marrow-derived macrophages were treated with or without IFNγ (1U/mL) for 24h and then infected with tachyzoites of the indicated strain for 24h and 48h. The cells were fixed and numbers of parasite/vacuole determined by immunofluorescence microscopy. Shown are the means ± SEM, pooled from three independent experiments; *ns= not significant*, \*\**P<*0.01, using a multiple comparison 2-way ANOVA test.

Next, mice were intraperitoneally infected with 500 tachyzoites of each strain and 30 days later brain cysts were enumerated. Our results showed that while cyst numbers were partially restored in mice infected with TgPHYaKO_II_:PHYa^PHYa^ parasites, brain cysts were undetectable in mice infected with up to 10^3^ TgPHYaKO_II_:PHYa^TUB^ parasites **(Fig. 7E)**.

### TgPHYa Is Important to Evade IFNγ-Dependent Nutritional Immunity

IFNγ kills *Toxoplasma* by upregulating the expression of proteins that degrade the parasitophorous vacuole membrane, increase reactive oxygen species, and limit essential nutrient (e.g. tryptophan) availability [28]. In response, *Toxoplasma* evades IFNγ effector mechanisms via the injection of parasite-encoded effector proteins that act to either prevent IFNγ from upregulating IFNγ-stimulated genes or inhibit the gene products’ activities [29]. To test whether TgPHYa interferes with parasite effector protein injection, we took advantage of the finding that TgIST is a secreted effector that inhibits IFNγ-stimulated gene expression and blocks IFNγ-dependent killing in host cells that are infected before exposure to IFNγ [30, 31]. HFFs were therefore infected with WT, TgPHYaKO_II_, TgPHYaKO_II_:PHYa^PHYa^, or TgPHYaKO_II_:PHYa^TUB^ tachyzoites and 24 h later treated with IFNγ or PBS as a vehicle control. The cells were fixed, and plaques numbers and sizes measured. We observed that loss of TgPHYa did not impact the ability for *Toxoplasma* to resist IFNγ-dependent killing under these conditions although we did note that IFNγ had a slight but reproducible effect on the size TgPHYaKO_II_:PHYa^PHYa^ plaques (**Figs. 8A and S5A**). We also observed that loss of TgPHYa did not increase parasite susceptibility to H_2_O_2_ (**Fig. 8B**). These data suggest that TgPHYa is dispensable for effector protein secretion or resistance to reactive O_2_ species.

Tryptophan is an essential amino acid for *Toxoplasma* and IFNγ limits available pools of tryptophan for parasites to scavenge as a result of upregulating the expression of the tryptophan catabolizing enzyme, indoleamine 2,3 dioxygenase (IDO) [19]. To test whether TgPHYa is important for tryptophan scavenging in IFNγ-treated cells, HFFs were stimulated with IFNγ for 24h (or PBS as a vehicle control) and then infected with WT, TgPHYa-deficient, or TgPHYaKO_II_ complemented tachyzoites. The cells were then grown in complete medium or medium supplemented with increasing levels of tryptophan. As expected, growth of all strains was prevented by IFNγ pretreatment (**Fig. 8C “+IFNγ” samples**). When the medium was supplemented with 16 or 32 µg/mL of tryptophan, the numbers of plaques formed were restored for all strains (**Fig. S5B**) as were the sizes of the plaques formed by WT and TgPHYaKO_II_:PHYa^PHYa^ parasites (**Fig. 8C**). In contrast, significantly smaller plaques were formed by knockout and TgPHYaKO_II_:PHYa^TUB^ parasites in tryptophan supplemented medium (**Fig. 8C**). To test if TgPHYa is also important for parasite growth under tryptophan restricted conditions in other cell types, we repeated these assays using murine bone marrow-derived macrophages and observed that only WT and TgPHYaKO_II_:PHYa^PHYa^ parasite growth was restored when exogenous tryptophan was added to IFNγ-treated BMDM (**Fig. 8D and Fig. S5E**).

The data that loss of TgPHYa, an enzyme localized to the parasite’s cytoplasm, does not impact effector secretion suggest that its role in survival under tryptophan depleted conditions is most likely in efficient uptake and/or utilization of the amino acid. We therefore hypothesized that virulence of TgPHYaKO_II_ parasites would be increased in IDO1 knockout mice since loss of IDO1 would result in increased levels of tryptophan available for parasites to scavenge. Thus, we assessed whether TgPHYa had an *in vivo* role in tryptophan utilization by comparing survival, weight loss and numbers of brain cysts of WT and IDO1^-/-^ mice infected IP with parental or TgPHYaKO_II_ parasites. While differences in survival were not observed in mice infected with low numbers of cysts (not shown), IDO^−/−^ mice infected with the TgPHYaKO_II_ had significantly increased morbidity as revealed by greater weight loss and increased brain cyst burdens in the IDO^-/^_-_ mice (**Fig. 9A-B**).

**Figure 9.**
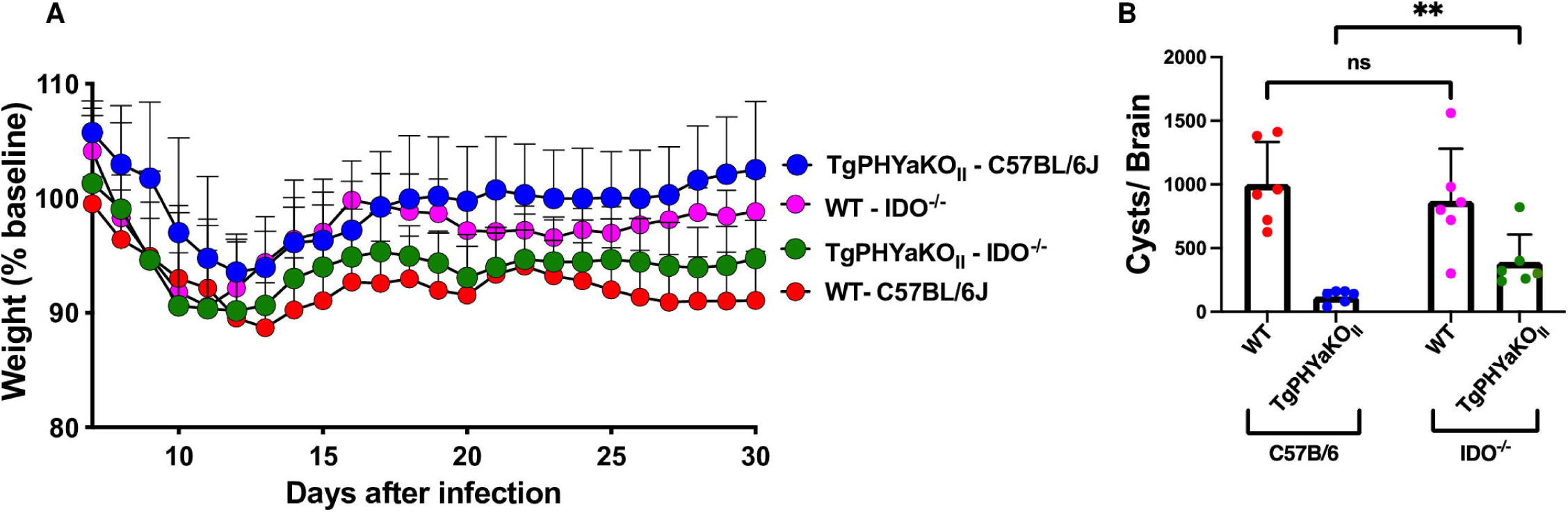
TgPHYaKO_II_ Is Hypervirulent in IDO1-Deficient Mice. **(A)** Kaplan-Meier survival curves of WT and IDO^-/-^ mice infected orally with 50 parental strain or TgPHYaKO_II_ cysts. Shown are cumulative data from 3 independent experiments (*n* = 9 total for each strain). **(B)** Brain cyst burdens in mice 30 days post infection. Shown are means ± SD (*ns= not significant*, \*\**P<*0.001, using a multiple Student’s *t* test with Holm-Sidake correction).

### Discussion

Hypoxic responses by pathogens and their hosts are important determinants for infection outcomes. Host immune responses are highly dependent on HIF-1-regulated transcription that is triggered by the hypoxic environment of infection foci [32–34]. Conversely, bacterial and fungal pathogens utilize O_2_ as a signaling molecule to regulate biofilm formation, virulence factor expression and host cell interactions [35–39]. In contrast to bacterial pathogens, less is known about the role of O_2_ sensing in protozoan virulence [4]. *Toxoplasma* expresses two oxygen sensing proteins - TgPHYb is important for growth at high O_2_ [40] and TgPHYa is important for growth at low O_2_ [15]. Yet the role that either PHD had in virulence was unknown. In this study, we generated a TgPHYa knockout mutant in the cystogenic type II ME49 strain background and demonstrated that it is important for virulence and brain cyst formation. While TgPHYaKO_II_ mutant parasites can establish an infection in the gut, they are unable to efficiently disseminate to peripheral tissues because TgPHYa is required to resist IFNγ-triggered killing and does so by overcoming IDO scavenging of tryptophan.

These data represent, to our knowledge, the first examples of an enzyme localized within the parasite cytoplasm that mediates resistance to IFNγ and of a PHD functioning as a virulence factor. One possible mechanism for how TgPHYa functions is that a proline in a tryptophan transporter is a TgPHYa substrate. But, we believe that is unlikely since growth defects are similar *in vitro* between parasites with knockout mutations in TgPHYa and TgGNT1, which is the first of a series of glycosyltransferases that modify prolyl hydroxylated TgSKP1 to generate a novel pentasaccharide knockout parasites [41]. Addition of this pentasaccharide is thought to alter the F-box protein repertoire that associates with the SCF-E3, which would impact SCF-E3-dependent protein ubiquitination [42]. Thus, we favor a model where TgPHYa alters SCF-E3-dependent ubiquitination of a tryptophan transporter or a transporter-associated protein that impacts the transporter’s stability, localization, and/or activity.

We found that TgPHYa overexpression has a dominant negative effect on IFNγ-controlled parasite virulence during the acute phase of the infection. This phenotype is reminiscent of the dominant negative effect that DdPHYa has on O_2_-dependent culmination during *Dictyostelium* development [43]. Since SKP1 was fully hydroxylated in both TgPHYaKO_II_:PHYa^PHYa^ and TgPHYaKO_II_:PHYa^TUB^ parasites, we do not believe that overexpression impacts TgPHYa activity. Rather, TgPHYa overexpression likely leads to aberrant complex formation that acts to sequester TgPHYa substrates and/or interacting proteins such as SKP1. Regardless of mechanism, the finding that TgPHYaKO_II_:PHYa^TUB^ parasites were virulent in IFNγ^-/-^ mice was surprising given its highly attenuated growth in tissue culture and avirulence in wild-type mice. This likely is due to the diverse repertoire *Toxoplasma* virulence factors that target distinct host resistance pathways [29].

We also observed that regardless of IFNγ expression growth of wild-type parasites was greater than that of TgPHYa knockout parasite growth. This could be due to TgPHYa also contributing to evasion of IFNγ-independent host resistance mechanisms such as TNFα [44] and IL-6 [45, 46]. Alternatively, TgPHYa may be important for the efficient utilization of other essential nutrients such as arginine. These hypotheses are not necessarily mutually exclusive since inducible nitric oxide synthase (iNOS), whose expression is regulated by cytokines such as IFNγ, TNFα, and IL-6 [45, 47], consumes arginine as a substrate. Earlier work by Denkers and colleagues demonstrated that the *Toxoplasma* virulence factor ROP16 impacts parasite growth by regulating iNOS expression [48].

The role of IDO in host resistance to *Toxoplasma* has been enigmatic. Early studies revealed that IFNγ-dependent induction of IDO1 expression inhibited parasite growth in human fibroblasts [19, 49, 50]. In addition, pharmacological inhibition of IDO enzymatic activity leads to increased mortality in *Toxoplasma*-infected mice [21]. But, IDO1 knockout mice did not display increased susceptibility to *Toxoplasma* infection and this is likely due to functional redundancy with a second IDO isoform, IDO2 [21, 51] or intrinsic differences in IDO enzymatic activity between mice and humans [52]. Our data that TgPHYa knockout parasites are more virulent in IDO1-deficient mice suggests that IDO1 is activated in parasite-infected mice but that it is unable to degrade tryptophan to levels sufficient to inhibit parasite growth. Since TgPHYa is an intracellular enzyme and that TgPHYaKO_II_ parasites have no apparent defects in secretion of parasite effectors into the host cell, we hypothesize that loss of TgPHYa leads a defect in tryptophan metabolism that is exacerbated under conditions of limited tryptophan availability. This is reminiscent of increased virulence of IDO^−/−^ mice infected with *Francisella tularensis* mutants with defects in tryptophan biosynthesis [53]. As discussed above, we hypothesize that loss of TgPHYa leads to defects in tryptophan uptake although we cannot rule out other pathways such as tryptophan catabolism or inefficient tRNA charging. Our future work will address these questions.

In summary, our data reveal a novel role for a pathogen-encoded cytosolic enzyme mediating resistance to IFNγ – tryptophan utilization efficiency. They also suggest that dual inhibition of host IDO and parasite tryptophan utilization represents a novel drug target against *Toxoplasma* and other tryptophan auxotrophic intracellular pathogens such as *Chlamydia* spp and *Trypanosoma cruzi* [54, 55].

## MATERIALS AND METHODS

### Cells and Parasites

The *Toxoplasma gondii* type II strain ME49 expressing red fluorescent protein (Table 1) and other strains generated here were maintained by passage in human foreskin fibroblasts (HFFs) (from American Type Tissue Culture, Reston, VA) in complete medium (Dulbecco’s modified Eagle medium supplemented with 10% heat-inactivated fetal calf serum, 1% L-glutamine, and 1% penicillin-streptomycin) as previously described [41].

### Disruption and Complementation of Tg*PHYa*

To generate TgPHYaKO_II_ the Skp1 prolyl 4-hydrohylase A gene (*PHYA*, TGME49_232960, Toxodb.org) was disrupted by a single-CRISPR/Cas9 method as previously described [56] with minor modifications. A single guide (sg) CRISPR plasmid, pSAG1::CAS9-U6::sgPHYa, was generated by site-directed mutagenesis of pSAG1::CAS9-U6::sgUPRT (a gift from Dr. David Sibley), using the Q5 site-directed mutagenesis kit from New England Biolabs (Ipswich, MA). The sgUPRT sequence was substituted by PCR with a guide DNA sequence targeting the catalytic domain of *PHYA* (embedded within primer A) and reverse primer B oriented in the opposite direction (Table S1). ME49-RFP tachyzoites (1.1 × 10^7^ in 400 µl) were co-transfected with pSAG1::CAS9-U6::sgPHYa (12.5 µg) and a dihydroxyfolate reductase (*DHFR*) amplicon (1 µg) [57]. TgPHYaKO_II_ parasites were subsequently selected in 1 µM pyrimethamine (Sigma) and insertion of the *DHFR* cassette was confirmed by PCR (Fig 1) using primers listed in Table S1.

To complement TgPHYaKO_II_ parasites, a *PHYA*-3×HA complementation plasmid with TgPHYa expressed under the *Toxoplasma* tubulin (TgTUB) promoter, was generated by modifying the UPRT Vha1 cDNA shuttle vector containing TgVhaI cDNA [58] using the NEB HiFi Builder method. The vector backbone, containing 973 nt of 5’- and 993 nt of 3’-*UPRT* flanking sequences, TgTUB promoter, and a 3×HA (C-terminal) sequence, was PCR amplified from the shuttle vector using primers G and H. The coding sequence of *TgPHYa* was PCR amplified from pET15b-TgphyA [15] using primers I and J. PCR fragments were gel-purified and incubated with HiFi DNA assembly enzyme mix (NEB) and transfected into *E. coli* Top10 cells (Thermofisher) to yield pUPRTPHYaKO_II_*:TgPHYa^TUB^* and sequence verified using primers I and J. TgPHYaKO_II_ parasites were transfected by co-electroporation with a PCR amplicon (generated using primers U and V) from pUPRTPHYaKO_II_*:TgPHYa^TUB^* (1 µg) and pSAG1::CAS9-U6::sgUPRT (10 µg), whose guide RNA targets the *UPRT* locus (Fig. S1B). Transfectants were selected in the presence of 10 µM fluorodeoxyuridine (FUdR, Sigma), and drug resistant clones were screened by PCR with primers listed in Table S1.

A second complementation construct was generated in which TgPHYa is expressed under the control of 1 kb of sequence upstream of the TgPHYa start codon (pUPRTPHYaKO_II_:*TgPHYa^PHYa^*). The upstream *TgPHYa* DNA fragment was generated by PCR of genomic DNA from ME49-RFP parasites using primers K and L, and the vector DNA fragment was generated from pUPRTPHYaKO_II_*:TgPHYa^TUB^* using primers M and N, which excluded the tubulin promoter sequence. Transfection, selection and validation of were performed as described above.

### Western Blotting

Tachyzoites were harvested from HFFs, counted with a hemacytometer, pelleted by centrifugation at 2,000 x *g* for 8 min at 4°C, and lysed in boiling SDS-PAGE sample buffer containing 5% 2-mercaptoethanol. Lysates from equivalent cell numbers were separated on a 4–12% gradient SDS-PAGE gel (Invitrogen;

Carlsbad, CA), transferred to nitrocellulose membranes that were blocked with 5% milk in TBS, and incubated with primary and secondary antibodies (Table 1). Blots were visualized and analyzed using a Li-Cor Odyssey scanner and software (Li-Cor; Lincoln, NE).

For SKP1 Western blots, extracts were prepared by vortexing parasites in 1% SDS, 10 mM Tris-HCl (pH 8.0), 1 mM EDTA, boiling for 5’, and centrifugation for 15’ at 21,000 × g. For reduction and alkylation [59], the supernatant was diluted with 4× sample buffer (0.9 M Tris (pH 8.45), 24% glycerol, 8% SDS, 0.01% phenol red, 0.01% Coomassie G) to which DTT was added (10 mM final concentration) and incubated at 65°C for 30’. Iodoacetamide was then added to a final concentration of 25 mM and incubated at room temperature in the dark for 1 h. Samples were electrophoretically separated on an Invitrogen NuPAGE 4-12% Bis-Tris gel (ThermoFisher) in MES-SDS buffer, pH 7.3 (ThermoFisher), transferred to nitrocellulose membrane, and Western blotted as above.

### Parasite Growth Assays

Plaquing assays were performed essentially as described [41]. Briefly, tachyzoites were added to confluent to HFF monolayers and grown undisturbed for 12 days. The cells were fixed with ice-cold methanol, stained with crystal violet, and plaques counted. Plaque areas were measured using ImageJ software (https://imagej.nih.gov/ij/). When assessing IFNγ/tryptophan supplementation, HFFs were pretreated for 24 h with 1 U/ml IFNγ (Sigma; St. Louis, MO) in media prepared using amino acid free DMEM supplemented, 3% dialyzed heat-inactivated fetal bovine serum, individual essential amino acids, and indicated concentrations of tryptophan. In vitro bradyzoite development was assessed as described [60]. Briefly, parasites were added to confluent HFFs seeded on glass coverslips in pH 8.1 Medium and grown for 72 h at 37°C at ambient CO_2_. Coverslips were fixed and then cysts were detected using Dolichos Biflorus lectin (Vector Lab, Burlingame, CA) and tachyzoites with anti-SAG1 antisera.

To assess growth in murine BMDMs, bone marrow cells isolated from the tibia and femur of C57BL/6 J mice were cultured in DMEM, 10% fetal bovine serum (FBS), 1% L-glutamine, 1% penicillin/streptomycin and 20% L929 conditioned medium for 5–7 days. BMDMs were then grown on coverslips in 24 well plates and stimulated with 1U/mL murine IFNγ (PeproTech; Cranbury, NJ, Cat#315-05) for 24 h pre infection with or without tryptophan as indicated. Both naïve BMDMs and IFNγ-activated BMDMs were infected at an MOI of 1. After 24 or 48 h, coverslips were fixed with 4% paraformaldehyde in PBS for 20 min at room temperature, blocked in 5% bovine serum albumin in PBS for 60 min, parasites detected by immunofluorescence staining with anti-SAG1 antibodies and Alexa Fluor 594-conjugated secondary antibodies. Coverslips were mounted with DAPI-containing VECTASHIELD mounting medium (Vector Labs; Burlingame, CA). At least 50 randomly selected vacuoles from 3 independent experiments were counted.

### Mouse Infections

Animal protocols and procedures were approved by the IACUCs at the University at Buffalo’s and University of Wyoming and carried out in accordance with Public Health Service policy on the humane care and use of laboratory animals and AAALAC accreditation guidelines. All mice were on a C57BL/6 background. C57BL/6 wild-type, IFN-γ^−/−^, CCR2^−/−^ and IDO^-/-^ were purchased from The Jackson Laboratory (Jackson Laboratory, Bar Harbor, ME). Morbidity was assessed by daily weighing and mortality by death or moribund state. Tissue cysts were enumerated 30 days post infection by counting numbers RFP^+^ cysts within three 25 μl aliquots of brain homogenates.

### NK Cell Depletion

Mice were depleted of NK cells using anti-NK1.1 (BioXcell; Lebanon, NH). Mice were treated with 200 mi of sterile PBS (vehicle alone) or 200 ug of anti-NK1.1 in 200 µl of sterile PBS intraperitoneally at day -1 of infection. Mice were injected again with vehicle or anti-NK1.1 at the same dose in the same volume at day 0 just prior to infection with either WT or TgPHYaKO_II_ tachyzoites. Mice were continually injected every other day with vehicle or anti-NK1.1 to maintain NK cell depletion during the course of the experiment.

### Quantitative PCR

Mice were sacrificed, tissues harvested and weighed, and genomic DNA isolated using the DNA Tissue kit (Omega, Norcross, GA). DNA (100 ng) was analyzed by real-time qPCR as described previously [40] using *Toxoplasma* B1 primers (*Forward:* 5′-TCCCCTCTGCTGGCGAAAAGT-3′, *Reverse:* 5′-AGCGTTCGTGGTCAACTA TCGATTG - 3′). Numbers of parasites were calculated from a standard curve generated in parallel using purified genomic DNA from known numbers of parasites. mRNA was analyzed by qRT-PCR as previously described [61] using the following *Toxoplasma*gene-specific primers: ß-actin - Forward: 5′- GCGCGACATCAAGGAGAAGC-3′, Reverse: 5′-CATCGGGCAATTCATAGGAC - 3′; and TgPHYa - Forward: 5′-TCTGTGGACACCAGAACTCG -3′, Reverse: 5′-CATGCAGGGAGTTCGCTGAG - 3′.

### Histology

Mice were euthanized and small intestines were removed and sections of the duodenum, jejunum and ileum immediately fixed in a 10% formalin solution. Paraffin-embedded sections were cut longitudinally at 0.5 μm and hematoxylin and eosin stained. Inflammation was scored in a in a blinded manner using a 0-4 scoring scale: 0, no inflammation; 1, slight infiltrating cells in lamina propria (LP) with focal acute infiltration; 2, mild numbers of infiltrating cells in the LP with increased blood flow and edema; 3, diffuse and massive infiltrating cells leading to disturbed mucosal architecture; 4, crypt abscess and necrosis of the intestinal villi. For each animal at least two sections (at least 100 μm from each other) were evaluated.

### Flow Cytometry

Small intestines without Peyer patches were harvested, minced, and incubated in digestion media (RPMI 1640, 1% penicillin-streptomycin, 1% l-glutamine, 0.1% beta-mercaptoethanol, 25 mM HEPES pH 7.0, 150 μg/ml DNase, and 59 μg/ml Liberase TL). Homogenates were passed through a 70-μm filter, washed in PBS, centrifuged at 1500 × *g* for 5 min, and cells were resuspended in RPMI 1640 supplemented with 10% FBS, 1% penicillin-streptomycin, 0.1% β-ME, 25 mM HEPES pH 7.0. Cells were stained with Live/Dead stain (ThermoFisher, Waltham, MA) or with antibodies against cell surface markers (Table 1). Peritoneal cells were similarly stained and harvested as described [60]. Samples were washed and resuspended in fluorescence-activated cell sorter (FACS) buffer (0.5 mM EDTA, 5% FBS, 0.001% sodium azide, and 1× PBS). Data were acquired using the BD LSR Fortessa cell analyzer and analyzed using FlowJo version 10.0.8 (TreeStar, Ashland, OR)

### Immunoprecipitation

Tachyzoites grown in HFF were harvested by syringe lysis and washed in ice-cold PBS. Total protein (1mg) was resuspended in 50 mM Tris-HCl pH 7.4 with 1% Triton X-100, 100 mM NaCl, 1 mM NaF, 0.5 mM EDTA, 0.2 mM Na3VO4, 1X protease inhibitor cocktail (Thermo Fisher Scientific), incubated on ice for 30 min and then subjected to sonication. Lysates were clarified by centrifugation at 16,000 xg, incubated with rabbit α-anti-HA clone C29F4 (Cell Signaling Technology) conjugated with protein G beads (Sigma-Aldrich) for 16 hours at 4 °C. Immune complexes were separated with SDS-PAGE and then Western blotted with antibodies against mouse anti-HA clone 6E2 (Cell Signaling Technology) or rabbit anti-TgSKP1 UOK75 [19].

### Statistics

All statistical assays were performed using GraphPad Prism v9.0 (GraphPad, La Jolla, Ca.).

## ACKNOWLEDGEMENT

We thank members of the Blader and West labs for their helpful discussions.

## FIGURE LEGENDS

**Supplemental Figure S1. CCR2^-/-^ Mice are Resistant to TgPHYaKO_II_ Infection.** C57BL/6J WT and C57BL/6J CCR2^-/-^ mice were orally infected by gavage with 50 cysts of ME49 WT or TgPHYaKO_II_ parasites. **(A)** Kaplan-Meier curve showing survival of C57BL/6J WT and C57BL/6J CCR2^-/-^ mice. Cumulative data from 3 independent experiments (*n* = 9 total for each strain). **(B)** Percent of weight loss of infected mice compared with initial weight before infection is plotted as mean ± SD from 3 independent experiments. **(C)** Cysts burden in surviving mice 30 days after infection is plotted as mean ± SD. The black line indicates the mean value. **(D)** C57BL/6J WT and C57BL/6J CCR2^-/-^ mice were orally infected by gavage with 50 cysts of ME49 WT or TgPHYaKO_II_ parasites. After 7 days infection, mice were euthanized and cells from small intestine lamina propria were processed and total number of infected cells was analyzed by flow cytometry as previously described (mean ± SEM, n = 6, pooled from three independent experiments). **(E)** IFN-γ levels in serum of C57BL/6J WT, IFN-γ ^-/-^ and CCR2^-/-^ mice 7 days after oral infection with 50 cysts of ME49 WT or TgPHYaKO_II_ parasites was quantified by an ELISA. (mean ± SEM, n = 2-6, pooled from three independent experiments).

**Supplemental Figure S2. NK cells Are Required for Early Control of WT and TgPHYaKO_II_.** Sham or NK-depleted mice were mock or parasite-infected intraperitoneally with 10^4^ WT or TgPHYaKO_II_ tachyzoites. After 5 days, peritoneal exudate was harvested and numbers of RFP^+^-infected cells were determined by flow cytometry. Shown are means ± SD of 2 independent experiments with 2 mice/experiment.

**Supplemental Figure S3. TgPHYa Complementation in TgPHYaKO_II_. (A)** Scheme for using CRISPR to target TgPHYa expression constructs to the UPRT locus. **(B).** Genomic DNA from the indicated strains were analyzed by PCR using the primers depicted in **(A)**.

**Supplemental Figure S4. TgPHYaKO_II_:PHYa^TUB^ Virulence is Highly Attenuated**. C57BL/6J WT mice were intraperitoneally infected with 10^6^ WT, TgPHYaKO_II_ or TgPHYaKO_II_:PHYa^TUB^ tachyzoites. **(A)** Kaplan-Meier curve showing survival of infected mice. Cumulative data from 3 independent experiments (*n* = 9 total for each strain). **(B)** Total number of parasite-infected peritoneal cells were determined by flow cytometry. Shown are mean ± SEM, n = 6, pooled from three independent experiments; \*\*\**P<*0.001, using unpaired Student *t* test).

**Supplemental Figure S5. Enumeration and Visualization Of the Effect of IFNγ On TgPHYa Knockout And Complemented Strains. (A)** HFFs were infected with *Toxoplasma* for 24 and then mock- or IFNγ-treated (1 U/ml). After 12 days, monolayers were fixed and numbers of plaques determined (mean ± SEM, pooled from three independent experiments). **(B)** HFFs were pretreated with IFNγ (1U/mL) for 24h and then infected with tachyzoites of the indicated strain. The cells were grown for 12 days in the presence of increasing tryptophan concentrations and then monolayers were fixed and plaque numbers enumerated. **(C)** Bone marrow-derived macrophages were treated with or without IFNγ (1U/mL) for 24h and then infected with tachyzoites of the indicated strain for 24h and 48h ± exogenous tryptophan. The cells were fixed and numbers of parasite/vacuole determined by immunofluorescence microscopy (Blue= DAPI, Red= SAG1). Data quantification are shown in Figure 8D.

**Table S1. List of Antibodies Used for This Study**

