## Supplementary material for "The *Toxoplasma* Oxygen Sensing Protein, TgPhyA, Is Required for Resistance to Interferon-gamma Mediated Nutritional Immunity": Fig S1

### Slide 1
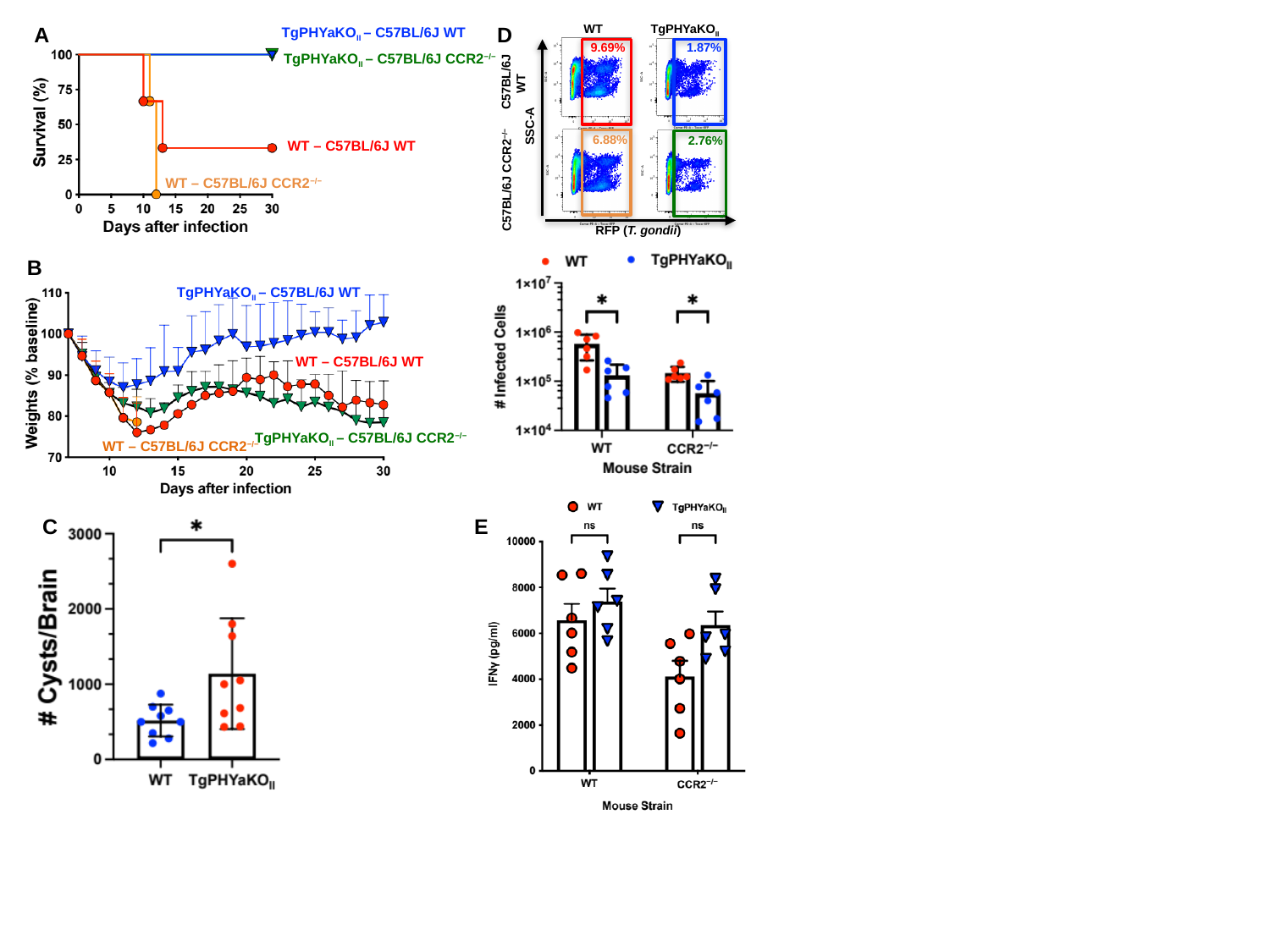

A
TgPHYaKO_II_ – C57BL/6J WT
TgPHYaKO_II_ – C57BL/6J CCR2^−/−^
WT – C57BL/6J WT
WT – C57BL/6J CCR2^−/−^
WT
TgPHYaKO_II_
9.69%
1.87%
C57BL/6J
WT
SSC-A
6.88%
2.76%
C57BL/6J CCR2^−/−^
RFP (T. gondii)
D
B
TgPHYaKO_II_ – C57BL/6J WT
WT – C57BL/6J WT
TgPHYaKO_II_ – C57BL/6J CCR2^−/−^
WT – C57BL/6J CCR2^−/−^
E
C
