## Supplementary material for "The *Toxoplasma* Oxygen Sensing Protein, TgPhyA, Is Required for Resistance to Interferon-gamma Mediated Nutritional Immunity": Fig S2

### Slide 1
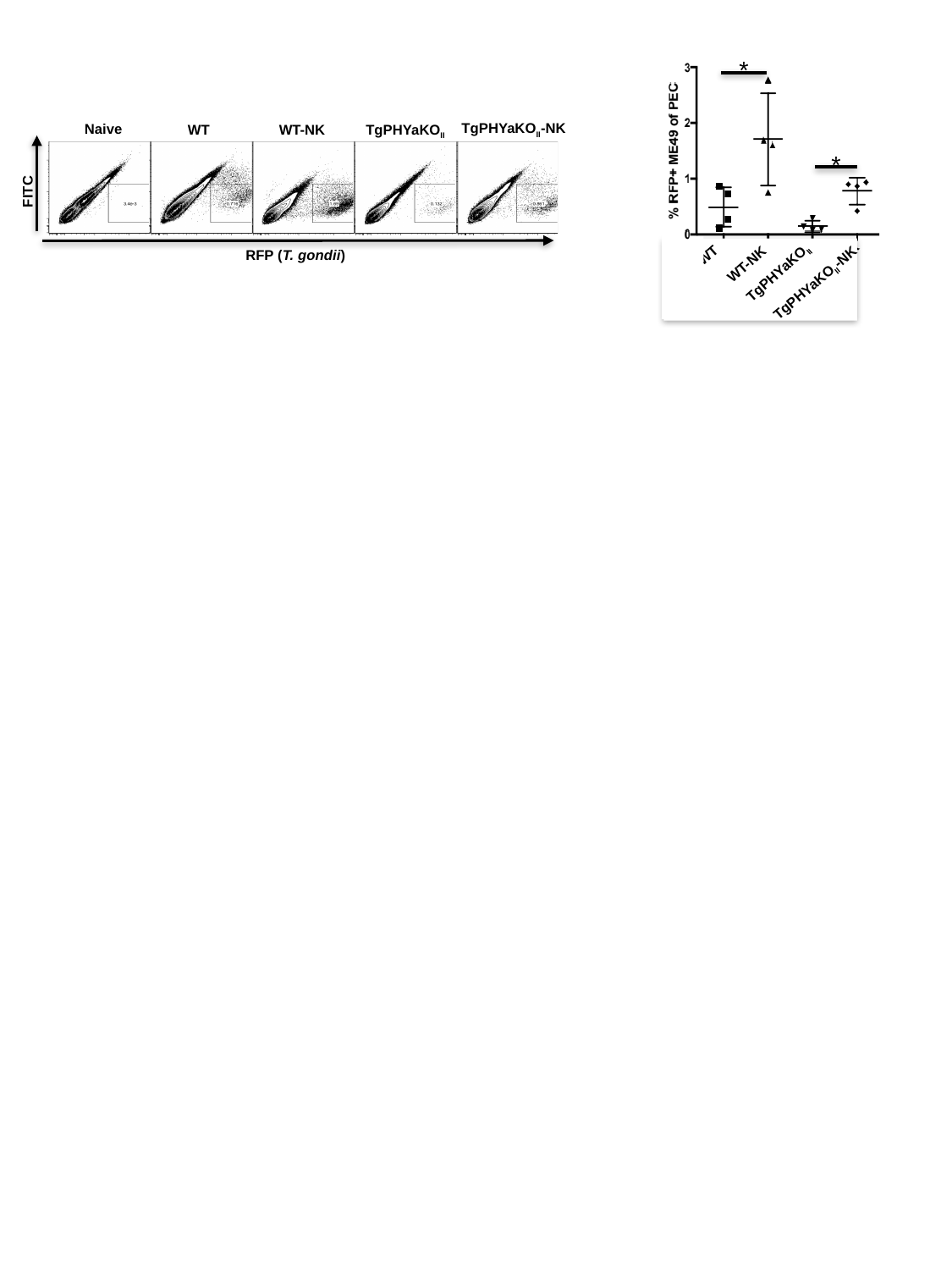

*
TgPHYaKO_II_-NK
Naive
TgPHYaKO_II_
WT
WT-NK
FITC
RFP (T. gondii)
*
FITC
WT
WT-NK
TgPHYaKO_II_-NK
TgPHYaKO_II_
