## Supplementary material for "The *Toxoplasma* Oxygen Sensing Protein, TgPhyA, Is Required for Resistance to Interferon-gamma Mediated Nutritional Immunity": Fig S3

### Slide 1
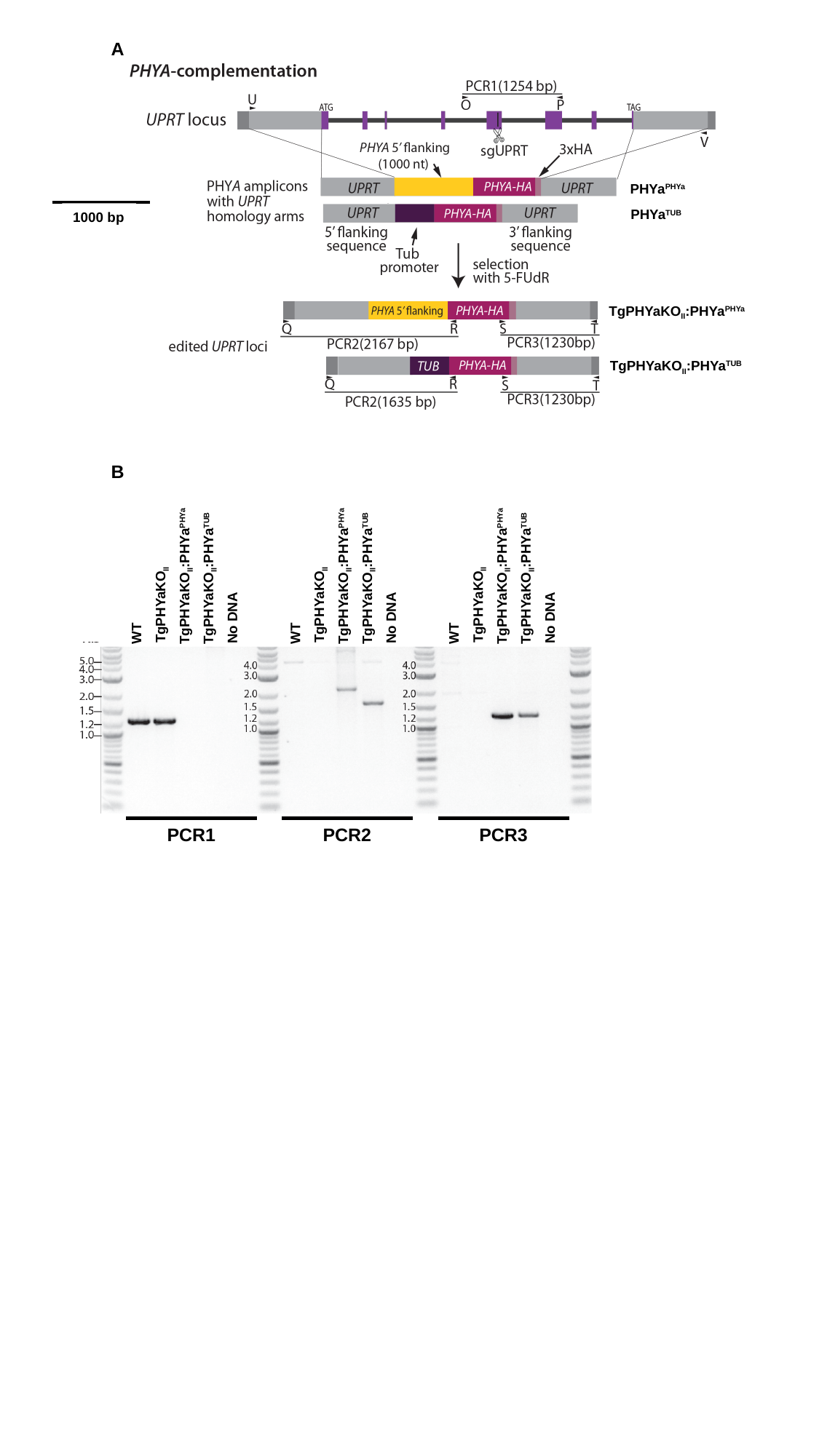

A
PHYa^PHYa^
PHYa^TUB^
1000 bp
TgPHYaKO_II_:PHYa^PHYa^
TgPHYaKO_II_:PHYa^TUB^
B
TgPHYaKO_II_:PHYa^PHYa^
TgPHYaKO_II_:PHYa^TUB^
TgPHYaKO_II_
No DNA
WT
TgPHYaKO_II_:PHYa^PHYa^
TgPHYaKO_II_:PHYa^TUB^
TgPHYaKO_II_
No DNA
WT
TgPHYaKO_II_:PHYa^PHYa^
TgPHYaKO_II_:PHYa^TUB^
TgPHYaKO_II_
No DNA
WT
PCR1
PCR2
PCR3
