## Supplementary figures and images for "The *Toxoplasma* Oxygen Sensing Protein, TgPhyA, Is Required for Resistance to Interferon-gamma Mediated Nutritional Immunity"

### Fig S4

#### Slide 1
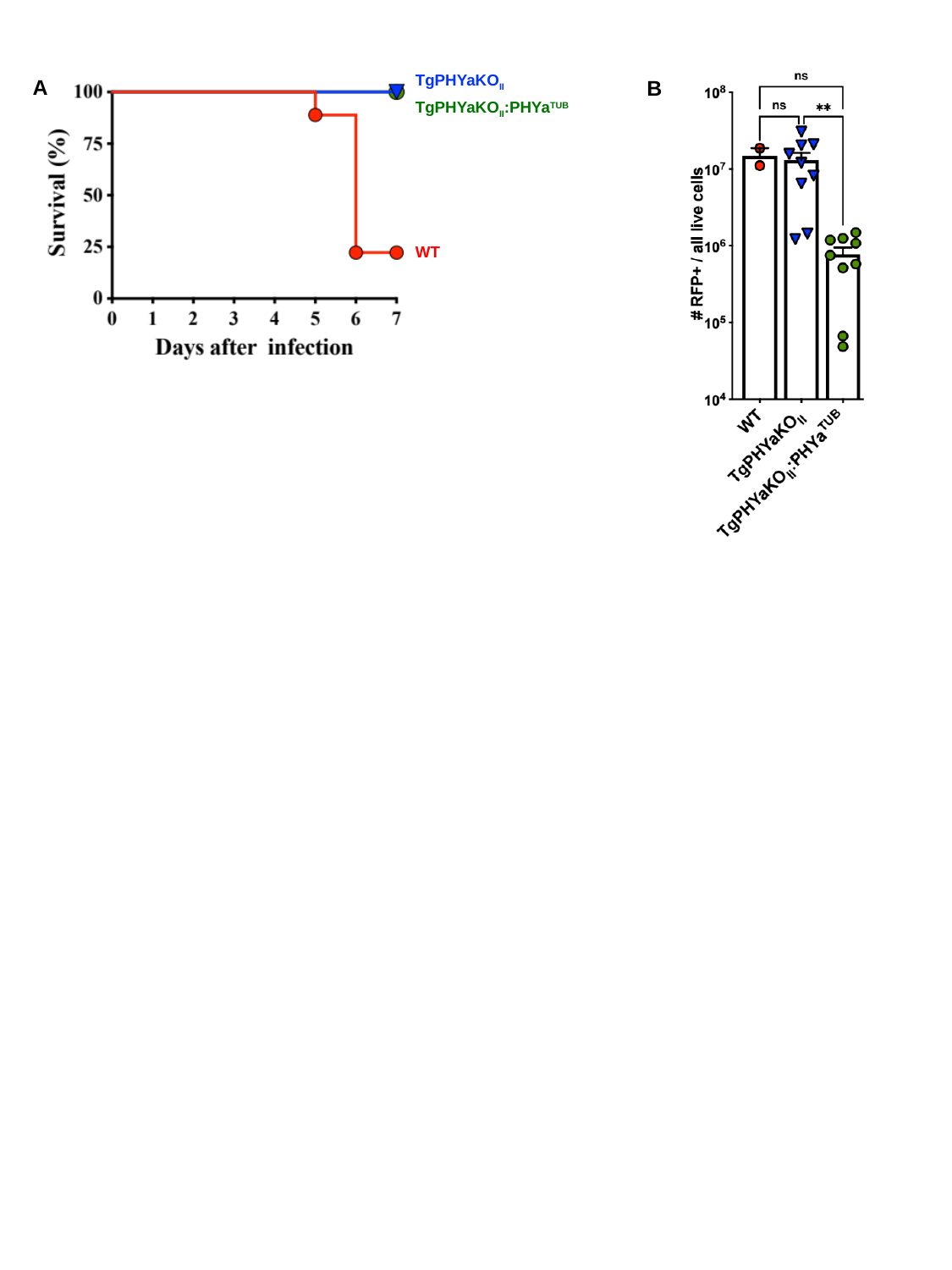

TgPHYaKO_II_
A
B
TgPHYaKO_II_:PHYa^TUB^
WT
