## Supplementary material for "The *Toxoplasma* Oxygen Sensing Protein, TgPhyA, Is Required for Resistance to Interferon-gamma Mediated Nutritional Immunity": Fig S5

### Slide 1
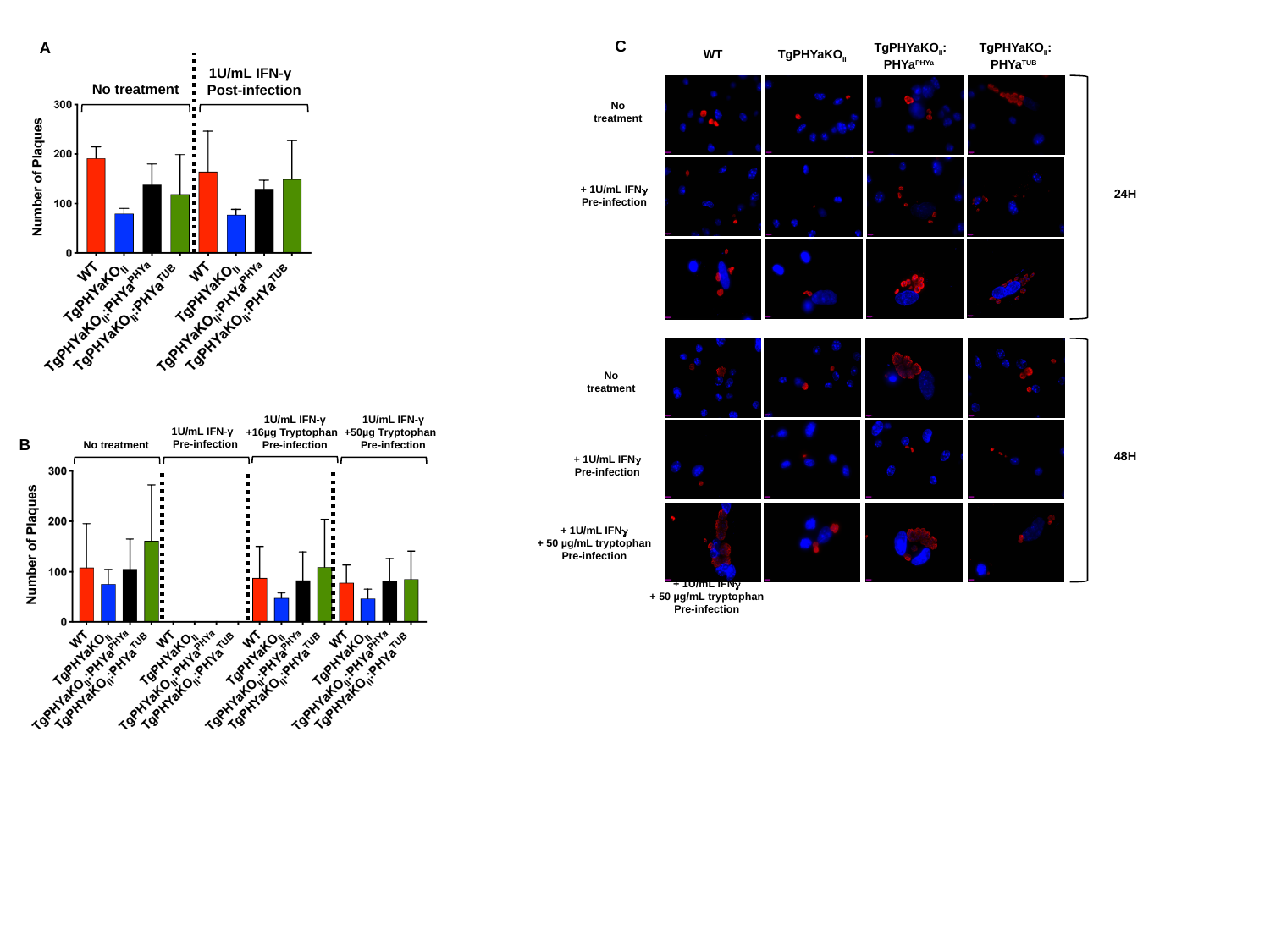

C
A
1U/mL IFN-γ
Post-infection
No treatment
TgPHYaKO_II_:PHYa^PHYa^
TgPHYaKO_II_:PHYa^TUB^
WT
TgPHYaKO_II_
No treatment
+ 1U/mL IFNγ
Pre-infection
24H
No treatment
48H
+ 1U/mL IFNγ
Pre-infection
+ 1U/mL IFNγ
+ 50 µg/mL tryptophan
Pre-infection
1U/mL IFN-γ
+16µg Tryptophan
Pre-infection
1U/mL IFN-γ
+50µg Tryptophan
Pre-infection
1U/mL IFN-γ
Pre-infection
No treatment
B
+ 1U/mL IFNγ
+ 50 µg/mL tryptophan
Pre-infection
